# Epigenetic age prediction using N6-methyladenine in the bumblebee *Bombus terrestris*

**DOI:** 10.1101/2025.06.20.660749

**Authors:** Thibaut Renard, Morgane Boseret, Serge Aron

## Abstract

Epigenetic alterations are a hallmark of aging. Age-specific DNA methylation patterns can be used to create ‘epigenetic clocks’—machine-learning algorithms that use methylation data from multiple genomic sites to predict an organism’s chronological age (*i*.*e*., the number of years or time passed since birth) or biological age (*i*.*e*., a measure of an organism’s health and functional status). Epigenetic clocks have been developed for mammals and, to a lesser extent, for birds, fish, amphibians, crustaceans, and insects. At present, all epigenetic clocks utilise C5-methylcytosine (5mC), a prevalent DNA methylation mark in vertebrates. However, in some species, 5mC marks are rare or even undetectable.

Here, we describe epigenetic clocks based on N6-methyladenine (6mA), a DNA methylation mark whose role in aging has remained unexplored. Using Oxford Nanopore Technology (ONT) sequencing, we measured genome-wide base-resolution levels of 6mA and 5mC in males of the buff-tailed bumblebee *Bombus terrestris* (*n* = 24). We constructed a series of epigenetic clocks using age-specific patterns in 6mA or 5mC. For each clock, predicted epigenetic age and chronological age were highly correlated. Furthermore, we pharmacologically increased individual lifespan with pharmacological agents and showed that, for individuals whose lifespan had been pharmacologically increased, each clock predicted younger epigenetic age than chronological age, indicating that the clocks captured signals of biological aging. Our results demonstrate that 6mA patterns can be used to build epigenetic clocks that accurately predict both chronological and biological age in animals, paving the way toward the use of 6mA as a reliable biomarker of aging.

## Introduction

Aging is a natural process during which molecular, cellular and physiological alterations accumulate in a time-dependent manner, resulting in compromised biological integrity and, consequently, increasing risk of intrinsic mortality (1–3). Epigenetic alterations—such as changes in DNA methylation, histone modifications, and chromatin structure—are a hallmark of aging (3); they dysregulate transcriptional profiles, impairing cellular and physiological homeostasis, leading to a decline of cellular and physiological integrity (4). The epigenome functions as a dynamic interface between genes and environmental signals, thereby enabling context-dependent regulation of gene expression (5). Due to its plasticity, the epigenome can be experimentally modulated to influence the aging process. Notably, interventions that restore youthful epigenetic patterns have been shown to re-establish gene expression profiles similarly to younger states, resulting in enhanced cellular function and effective biological rejuvenation in both mammalian cells and whole organisms (6–8).

DNA methylation (DNAm) is the most extensively studied epigenetic mark linked to the aging process (9, 10). During aging, the DNA methylome exhibits two contrasting types of alterations— stochastic *versus* predictable—both of which contribute to age-related phenotypes. Stochastic changes accumulate randomly across the genome over time, increasing epigenetic entropy and transcriptional noise, which can disrupt cellular function (4, 8). In contrast, predictable changes occur consistently at specific genomic loci and can be leveraged to build epigenetic age prediction models (epigenetic clocks) which quantitatively predict chronological age (the actual time since birth) and/or biological age (an estimate of biological function) (9, 11). Epigenetic clocks have been primarily developed in mammals (12) and, to a lesser extent, in amphibians (13), fishes (14), crustaceans (15), and insects (16). Notably, these clocks can be trained to predict age across different cell, tissues, and even species, indicating that some features of epigenetic aging are evolutionarily conserved (12). The epigenetic clock theory of aging proposes that DNA methylation age represents the molecular traces left by both developmental and maintenance programs, thereby capturing intrinsic aging processes that contribute to functional decline in tissues and are conserved across organisms (9). Because these age-related epigenetic changes are consistent across different species and levels of biological organization, from individual cells to whole organisms, epigenetic clocks offer powerful tools to identify evolutionary conserved molecular signatures of aging, which are more difficult to consistently identify at the transcriptomic and proteomic levels due to high context-dependence (17). Despite this advantage, the proximal mechanisms underlying the existence of epigenetic clocks and their evolutionary consequence remain incompletely understood. Recent findings indicate that epigenetic clocks correlate with some hallmarks of aging, but not all (18), and capture both aging-promoting and aging-buffering changes, suggesting they reflect a complex interplay between deterioration and adaptation (19). Importantly, all existing epigenetic clocks developed to date rely exclusively on C5-methylcytosine (5mC), the predominant form of DNA methylation in eukaryotes (20). However, in several taxa, 5mC is extremely low (*e*.*g*., bees (21) and fruit flies (22)) or even undetectable (*e*.*g*., nematodes (23) and yeasts (24)), raising the question of whether alternative epigenetic marks could serve as surrogates of aging biomarkers in these species.

N6-methyladenine (6mA) is a distinct form of DNAm, originally thought to be exclusive to prokaryotes where it plays a key role in cellular immunity (25, 26). Recently, 6mA has been identified in various eukaryotes—including *Chlamydomonas* (27), *C. elegans* (28), *D. melanogaster* (29), *X. laevis* (30), mice (30), and humans (30)—where it is involved in gene expression regulation. Here, we show that 6mA can serve as a reliable biomarker of both chronological and biological aging. Using long-read Oxford Nanopore Technologies (ONT) sequencing, we generated genome-wide, base-resolution levels of 6mA and 5mC across the DNA methylome of buff-tailed bumblebee (*Bombus terrestris*) males spanning their entire adult lifespan. First, we compared age-associated patterns in 6mA and 5mC using differential methylation analyses, methylation entropy, and rate of change quantification. Second, we used these age-related signatures to construct a series of epigenetic clocks based on either 6mA or 5mC, and found that both correlated with chronological age with similar accuracy. Third, we pharmacologically extended male lifespan and showed that both 6mA- and 5mC-based clocks reported reduced epigenetic age in treated males, indicating their ability to track biological aging. These findings highlight that 6mA can serve as an accurate proxy of chronological and biological age in *B. terrestris*, paving the way toward its use among animals.

## Results

### Aging DNA methylome

DNA methylation patterns experience conserved alterations with age, including global hypomethylation, site-specific hypermethylation, increased methylation variability (entropy), and linear shifts in methylation levels over time (11, 31, 32). These patterns have been extensively characterized for 5mC. To test whether they also occur for 6mA, we sequenced the entire DNA methylome of *B. terrestris* males using long-read Oxford Nanopore Technologies (ONT) sequencing to generate genome-wide, base-resolution levels of 6mA and 5mC in males of three discrete age groups (7, 21, or 35-day-old), spanning the entire lifespan of *B. terrestris* adult males.

First, we investigated whether shifts in global methylation levels occur for 6mA and/or 5mC by quantifying global, genome-wide levels of 6mA and 5mC. Global levels of 6mA were consistently higher than 5mC across the three age groups (mean percentage of methylated sites ± SD: 6mA = 2.85 ± 0.44 % and 5mC = 1.87 ± 0.83 %, Wilcoxon Mann-Whitney U test: *U* = 199, *p* = 1.4e-04). Neither methylation type experienced significant variation in global levels across male lifespan (mean global methylation levels ± SD: 6mA: 7-day-old males = 2.71 ± 0.33 %, 21-day-old males = 2.85 ± 0.55 %, 35-day-old males = 2.99 ± 0.48 %, Kruskal-Wallis test: *chi-squared* = 0.06, *df* = 2, *p* = 0.97, Figure 1A; 5mC: 7-day-old males = 1.72 ± 0.29%, 21-day-old males = 2.09 ± 1.39%, 35-day-old males = 1.79 ± 0.54 %, Kruskal-Wallis test: *chi-squared* = 0.14, *df* = 2, *p* = 0.93, Figure 1B), indicating that the DNA methylome does not undergo global hypomethylation in this species.

**Fig. 1.**
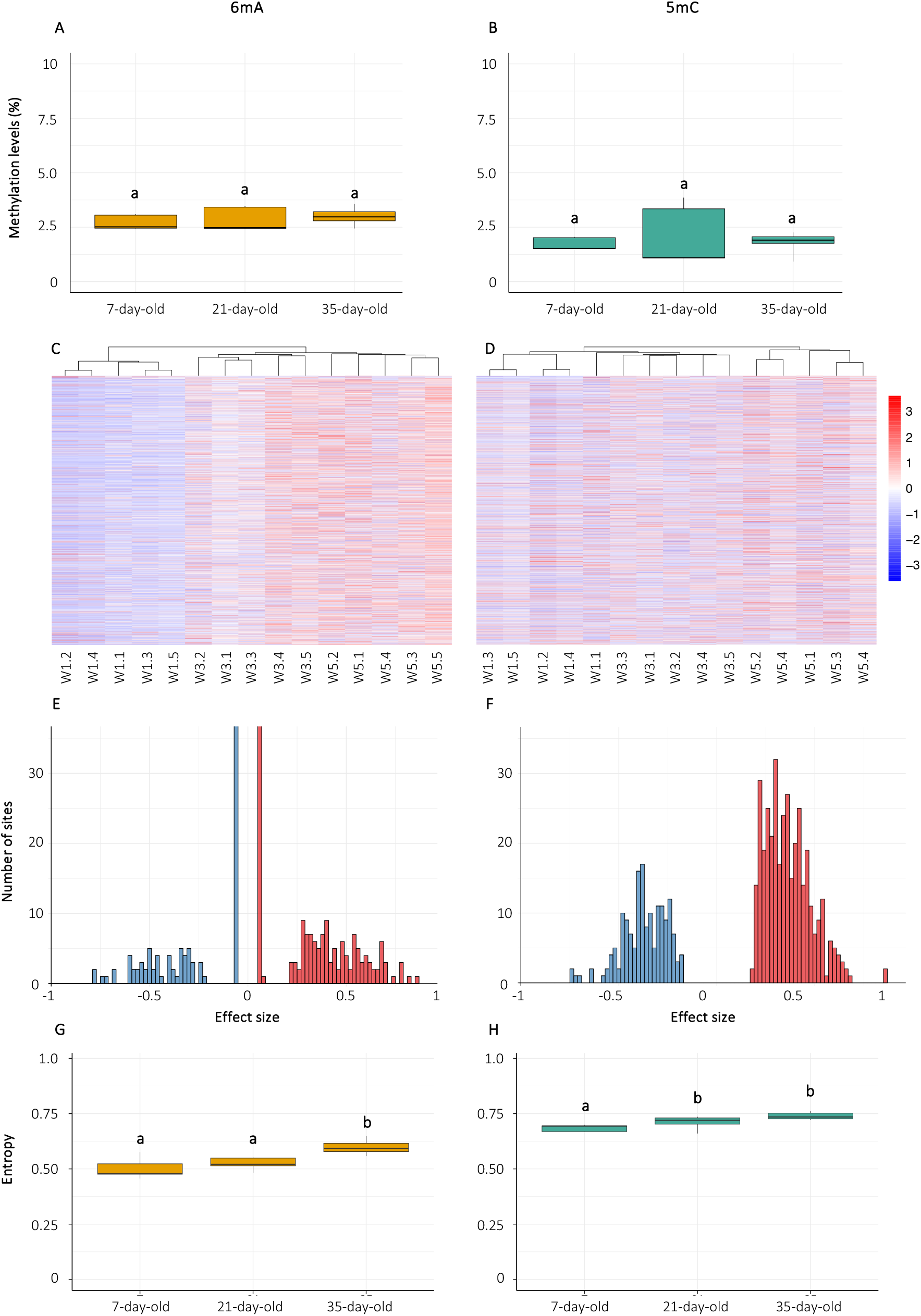
Age-related changes in 6mA and 5mC in *Bombus terrestris* males. **(A)** Global levels of 6mA and **(B)** 5mC measured across the genome in 7-day-old, 21-day-old, and 35-day-old males. **(C-D)** Heatmaps with hierarchical clustering showing base-resolution methylation levels of 6mA or 5mC at differentially methylated sites (DMS), respectively. Methylation levels were normalized per site using Z-score transformation to allow relative comparison between age groups (W1= 7-day-old, W3 = 21-day-old, W5 = 35-day-old). Blue indicates below-average and red indicates above-average methylation relative to the mean. **(E-F)** Distribution of effect sizes for 6mA and 5mC DMS between 35-day-old and 7-day-old (W1) males. Effect size is defined as the difference in mean methylation level between 35-day-old and 7-day-old individuals, such that positive values indicate hypermethylation and negative values indicate hypomethylation in older males. **(G-H)** Average methylation entropy at 6mA and 5mC DMS across age groups. Statistical significance is denoted by different letters above the boxplots (*p* < 0.05). *n* = 5 per age group.

Second, we explored if site-specific alterations occur within the DNA methylome throughout males’ lifespan by performing differential methylation analysis between 7-day-old *vs* 35-day-old. Only extreme age groups were considered to capture the strongest age-related alterations. Using a ready-to-implement Bayesian framework from ONT, we identified 850 differentially methylated sites (DMS) for 6mA and 530 DMS for 5mC. Hierarchical clustering based on 6mA or 5mC DMS grouped individuals of the same age together (Figures 1C-D), showing their methylomic similarities. The ratio of hypomethylated-to-hypermethylated sites differed between 6mA and 5mC: while most DMS for 6mA showed increased methylation with age, the proportion of 5mC that gained and lost methylation over time was more balanced (proportion of sites in 35-day-old *versus* 7-day-old: 6mA: 17.2 hypomethylated and 82.8 % hypermethylated, Figure 1E, 5mC: 30.9 % hypomethylated and 69.1 % hypermethylated, Figure 1F, chi-squared test: *chi-squared* = 34.736, *df* = 1, *p* = 3.775e-09). Several DMS for 6mA and 5mC were found within non-coding RNAs (ncRNA). Specifically, 17.0 % of 6mA DMS were found in ncRNA and 82.0 % in protein-coding genes. In contrast, only 2.6 % of 5mC DMS occurred in ncRNAs, with the vast majority (97.4%) located in protein-coding regions. This difference in distribution between 6mA and 5mC DMS was statistically significant (chi-squared test: *chi-squared* = 65.4, *df* = 1, *p* = 6.115e-16). In addition, ∼1% of 6mA DMS were found in ribosomal DNA (rDNA), whereas no 5mC DMS were detected in rDNA. The 6mA DMS with the greatest methylation difference was located in the *TfIIFbeta* [*transcription factor TFIIFbeta*] gene, which encodes a subunit of the TFIIF transcription factor complex—an essential component of the RNA polymerase II pre-initiation complex (33). The cytosine displaying the largest methylation change was located in *RAPK2* (*rho-associated protein kinase 2*), whose protein product is a serine-threonine kinase that regulates cell shape and movement by acting on the cytoskeleton (34). The complete list of 6mA and 5mC DMS can be found in ESM Tables S1-S2. Functional enrichment analyses using gene ontology (GO) terms were performed to identify the biological processes (BP) associated with the aging DNA methylome of *B. terrestris* males. We identified 58 and 54 significantly enriched BP GO terms for 6mA or 5mC DMS, respectively. Enriched BP terms associated with 6mA were involved in development, and epigenetic and epitranscriptomic regulation. Enriched 5mC BP terms were mostly linked to proteostasis. The complete list of GO terms can be found in ESM Tables S3-S4.

Third, we examined whether the *B. terrestris* DNA methylome exhibits stochastic age-related changes in 6mA and 5mC. To this end, Shannon entropy was calculated for each 6mA or 5mC across the genome of each sample, then averaged across sites within sample to derive an individual-level measure of methylomic entropy (11). Genome-wide methylomic entropy did not undergo significant increase with age for either methylation type (mean methylomic entropy ± SD: 6mA: 7-day-old = 0.32□±□0.02, 21-day-old = 0.34□±□0.03, 35-day-old = 0.34□±□0.03, one-way ANOVA: *F* = 1.014, *df* = 2, *p* = 0.386, 5mC: 7-day-old = 0.32□±□0.02, 21-day-old = 0.34□±□0.04, 35-day-old = 0.36□±□0.04, one-way ANOVA: *F* = 1.48, *df* = 2, *p* = .259). However, 6mA and 5mC methylomic entropy increased when considering DMS only (6mA: 7-day-old = 0.50□±□0.05, 21-day-old = 0.52□±□0.03, 35-day-old = 0.61□±□0.03, one-way ANOVA: *F* = 12.66, *df* = 2, *p* = 0.0006, post-hoc Tukey HSD: 7-day-old *vs* 35-day-old: *p* = 0.0009, 21-day-old *vs* 35-day-old: *p* = 0.0072, Figure 1G, 5mC: 7-day-old = 0.68□±□0.02, 21-day-old = 0.71□±□0.03, 35-day-old = 0.75□±□0.01, one-way ANOVA: *F* = 10.77, *df* = 2, *p* = 0.0013, post-hoc Tukey HSD: 7-day-old *vs* 35-day-old: *p* = 0.0010, Figure 1H).

### Epigenetic age prediction

Next, we investigated whether linear changes in 6mA and 5mC occur within the aging DNA methylome of *B. terrestris* and, if so, whether they can be leveraged to build epigenetic age predictors (epigenetic clocks).

We conducted a site-specific rate of change (ROC) analysis to identify linear changes in 6mA or 5mC. For each DMS, we applied simple linear regression models to quantify the relationship between methylation levels and chronological age and used the model slope as a measure of the site’s ROC. Mean ROC was low for both methylation types; however, 6mA ROC was significantly higher than 5mC ROC (mean ROC per week ± SD [number of significant sites retained]: 6mA = 1.32□±□2.04 % [*n*□=□473], 5mC = 0.61□±□11.14 % [*n*□=□282], Wilcoxon rank-sum test: *W* = 60.591, *p* = 0.035). These results show that linear age-related changes occur for both 6mA and 5mC. In addition, the mean rate of change (ROC) in 5mC methylation showed significantly greater variance compared to 6mA (Levene’s test for homogeneity of variance: *F* = 712.73, *df* = 1, *p* < 2.2e-16), suggesting that 5mC DMS exhibit a broader range of methylation dynamics than 6mA.

To explore whether linear changes in 6mA or 5mC levels can be leveraged to build epigenetic clocks, we applied penalized regression models to develop a series of clocks based on either 6mA or 5mC. We adopted a three-step sequential approach to fully explore and exploit different features of the aging DNA methylome. Each model was trained using a leave-one-out cross-validation (LOOCV) approach to minimize the risk of overfitting, then tested on an independent (testing) dataset.

First, we applied elastic net regressions on methylation levels (6mA or 5mC) to identify sites that correlate with chronological age. During model training, elastic net regressions selected 48 adenine and 44 cytosine residues whose methylation levels strongly correlate with chronological age (training dataset: 6mA clock: correlation = 0.999, mean relative error (MRE) = 2.6 %, mean absolute error (MAE) = 0.3 days, *p* (linear model) < 2e-16; 5mC clock: correlation = 0.998, MRE = 3.1 %, MAE = 0.5 days, *p* (linear model fitting) < 2e-16). Among these, 9 adenine and 40 cytosine residues were located within coding regions. For the 6mA clock, several sites were found near ribosomal DNA. Cytosines from the 5mC clock were not significantly enriched in any biological process. When applied to the testing dataset, both 6mA and 5mC elastic net models accurately predicted chronological age (testing dataset: 6mA clock: correlation = 0.987, mean relative error (MRE) = 7.3 %, mean absolute error (MAE) = 1.8 days, Figure 2A; 5mC clock: correlation = 0.978, MRE = 11.4 %, MAE = 2.6 days, Figure 2B), despite slight underestimation of predicted age for 21-day-old males for the 5mC clock.

**Fig. 2.**
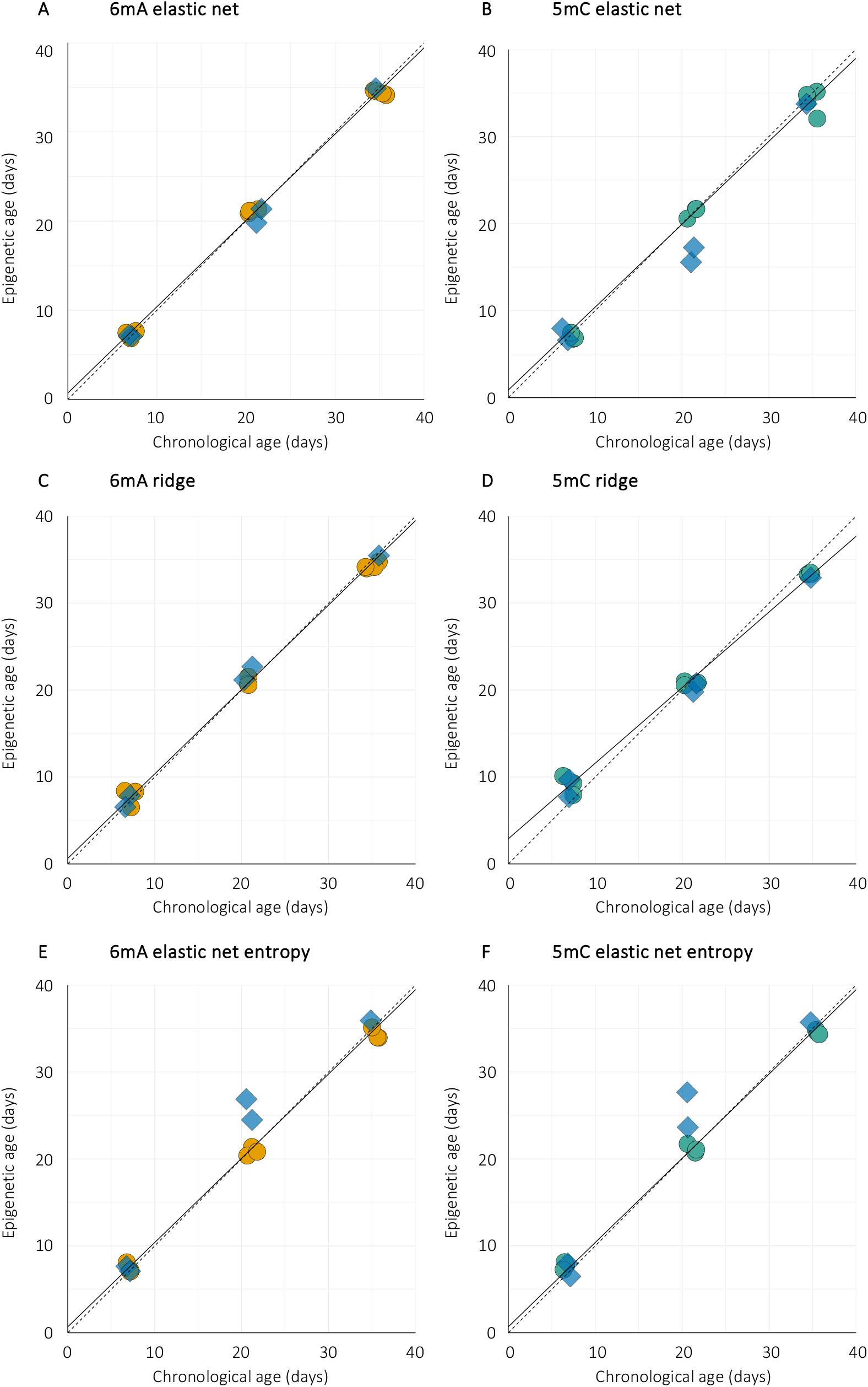
Epigenetic age prediction models in *Bombus terrestris* males. Epigenetic clocks were constructed using penalized regression models with leave-one-out cross-validation (LOOCV), based on age-associated patterns in 6mA **(A,C,E)** or 5mC **(B,D,F). (A–B)** Elastic net regression models were trained on genome-wide methylation levels. **(C–D)** Ridge regression models were trained on methylation levels of differentially methylated sites (DMS). **(E–F)** Elastic net regression models were trained on entropy values computed at each DMS. Solid lines indicate the fitted epigenetic age prediction models. Dashed lines represent the identity line (y = x), showing the expected relationship under perfect age prediction. Data points represent individual samples: orange (6mA) and green (5mC) circles correspond to training samples; blue diamonds represent test samples. *n* = 5 per age group.

Second, we investigated whether epigenetic clocks could be generated using DMS only. To this end, we used ridge regressions to build models using all 6mA or 5mC DMS as this type of penalized regression shrinks the coefficient of weakly predictive sites toward zero, without driving them entirely to zero. As expected, model training reported very high age prediction accuracy (training dataset: 6mA clock: correlation = 0.999, mean relative error (MRE) = 3.0 %, mean absolute error (MAE) = 0.4 days, *p* (linear model) < 2e-16; 5mC clock: correlation = 0.9993, MRE = 10.6 %, MAE = 1.3 days, *p* (linear model) = 6.9e-13). Both 6mA and 5mC ridge models accurately predicted age in the naïve dataset (testing dataset: 6mA clock: correlation = 0.985, mean relative error (MRE) = 5.4 %, mean absolute error (MAE) = 1.6 days, Figure 2C; 5mC clock: correlation = 0.9983, MRE = 10.7 %, MAE = 1.3 days, Figure 2D).

Finally, we tested whether site-specific methylation entropy could be used as a surrogate for methylation levels in epigenetic clocks. While 5mC entropy has recently been shown to correlate with chronological age (35), the potential use of 6mA entropy remains unknown. We used elastic net regression to select sites whose methylation entropy correlate with chronological age. Both entropy-based models reported accurate performance metrics (training dataset: 6mA clock: correlation = 0.9972, mean relative error (MRE) = 3.3 %, mean absolute error (MAE) = 0.6 days, *p* (linear model) < 2e-16; 5mC clock: correlation = 0.9995, MRE = 3.1 %, MAE = 0.5 days, *p* (linear model) < 2e-16; testing dataset: 6mA clock: correlation = 0.9528, mean relative error (MRE) = 11.4 %, mean absolute error (MAE) = 2.6 days, Figure 2E; 5mC clock: correlation = 0.9575, MRE = 10.4 %, MAE = 2.3 days, Figure 2F). Elastic net regressions selected 37 adenines and 25 cytosines whose entropy methylation best predicted chronological age. As for the 6mA clock based on methylation levels, several adenine residues from the 6mA entropy clock were located within or close to ribosomal DNA. Predictive cytosines from the 5mC entropy clock were found in genes involved in DNA replication, transcription and DNA damage repair. Both entropy clocks slightly overestimated age in 21-day-old males.

### Pharmacological lifespan extension and biological aging

Epigenetic clocks can be trained to predict either chronological or biological age. Our findings show that 6mA and 5mC correlate with chronological age in *B. terrestris* males; whether they are also associated with biological age remains to be tested. Since our experiments were conducted under standardized laboratory conditions, we did not expect significant inter-individual variation in biological age between individuals. Therefore, we used pharmacological agents to increase individual lifespan, thus affecting biological age, and we tested whether the 6mA- and 5mC-based clocks capture signals of biological aging.

First, we tested if bumblebee males’ lifespan can be pharmacologically extended by chronically feeding them with rapamycin and resveratrol, two well-researched agents which act upon the lifespan-regulation mTOR (mechanistic target of rapamycin) and sirtuins pathways, respectively. The mean lifespans of males fed rapamycin and resveratrol were increased by 37% and 34%, respectively, compared to males fed a control solution containing sugar syrup and DMSO (mean lifespan ± SD: control = 26.9 ± 7.4 days, rapamycin = 37.6 ± 8.8 days, resveratrol = 36.5 ± 7.4 days, Kruskal-Wallis: *chi-squared* = 42.388, *df* = 2, *p* = 6.2e-10, Figure 3A). Both treatments caused a 31% increase in maximum lifespan (control = 51 days; rapamycin = 67 days; resveratrol = 66 days) (Figure 3B).

**Figure 3.**
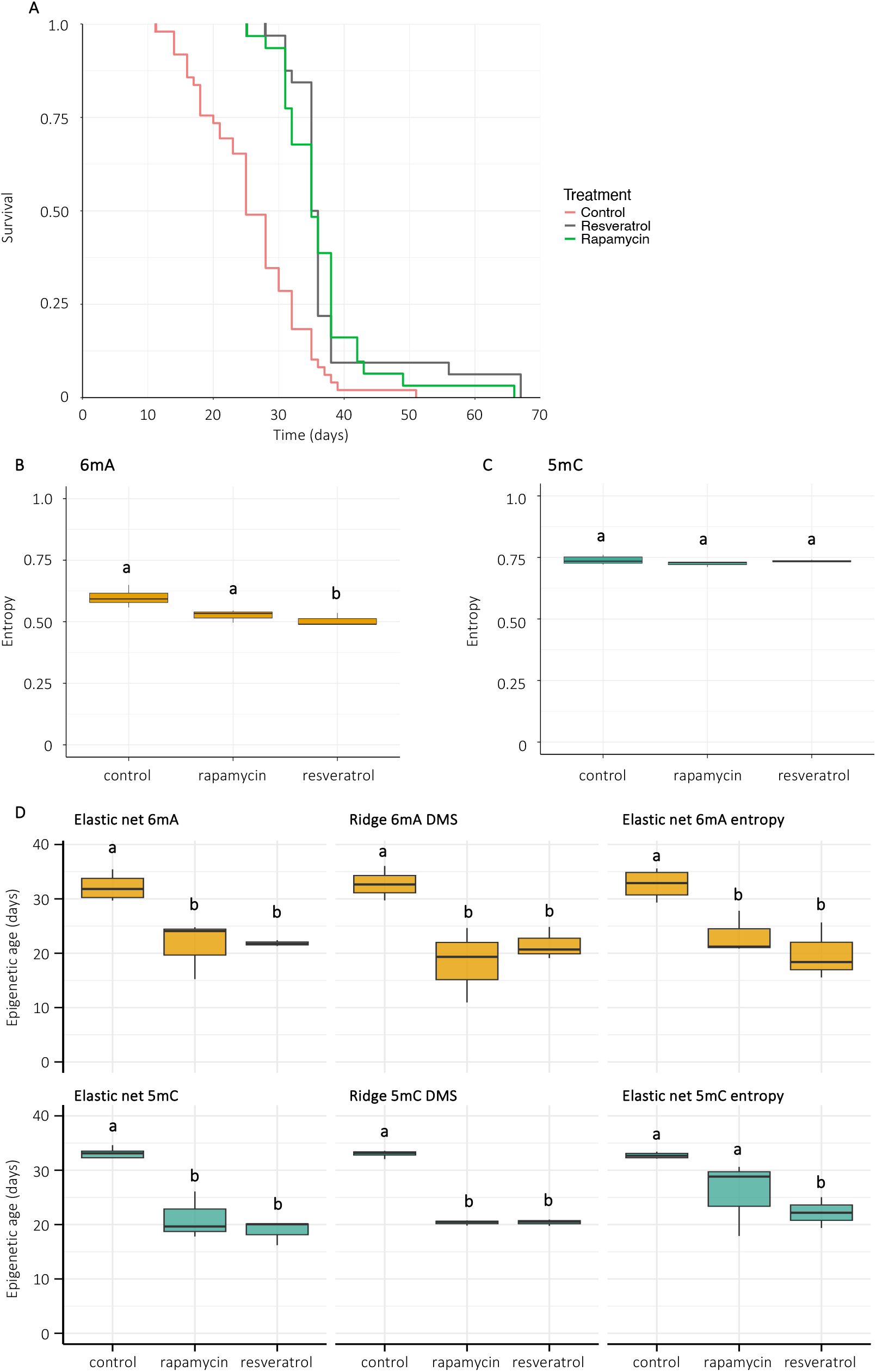
Pharmacological lifespan extension is associated with reduced epigenetic age in 35-day-old *B. terrestris* males. **(A)** Survival curves of males fed a control solution (light coral, *n* = 49), rapamycin (green, *n* = 32) or resveratrol (grey, *n* = 31) (Cox proportional hazards regression model: rapamycin-fed males: *p* = 6.61e-07, resveratrol-fed males: *p* = 8.98e-07). **(B-C)** Entropy levels of 6mA and 5mC DMS, respectively (*n* = 3 per treatment group). **(D)** Predicted epigenetic ages of 35-day-old males fed rapamycin or resveratrol (*n* = 3 per treatment group). Different letters above the box represent significant statistical differences (*p* < 0.05) using either one-way ANOVA and post-hoc Tukey HSD test or Kruskal-Wallis and post-hoc Wilcoxon-Mann-Whitney pairwise test.

Second, we investigated whether pharmacological lifespan extension is associated with methylomic changes. We sequenced the DNA methylome of 35-day-old rapamycin- or resveratrol-fed males and compared it to 35-day-old control males. Global DNAm levels did not vary significantly between control males, rapamycin- and resveratrol-fed males (6mA: control = 3.37 ± 0.21 %, rapamycin: 2.95 ± 0.42 %, resveratrol: 2.88 ± 0.31 %, one-way ANOVA: *F* = 1.972, *df* = 2, *p* = 0.22, 5mC: control = 1.84 ± 0.42 %, rapamycin = 2.08 ± 0.26 %, resveratrol = 2.07 ± 0.13 %, one-way ANOVA: *F* = 3.028, *df* = 2, *p* = 0.11). Differential methylation analyses revealed that both treatments had much stronger effects on 5mC than 6mA (6mA DMS: rapamycin = 10, resveratrol = 4; 5mC DMS: rapamycin = 14,745, resveratrol = 8,081). Both lifespan-extending treatments induced 5mC DMS, but not 6mA DMS, in genes related to their target protein (rapamycin: *raptor* [*regulatory associated protein of MTOR complex 1*], *lamtor* [*late endosomal/lysosomal adaptor, MAPK and MTOR activator*], resveratrol: *sirt1* [*NAD+-dependent protein deacetylase sirtuin-1*] (ESM Tables S5-S6). Functional enrichment analyses for 6mA identified 37 and 6 enriched BP terms between rapamycin- or resveratrol-fed *versus* control males. For 5mC DMS, 277 and 283 BP terms were enriched between rapamycin or resveratrol males *versus* the controls. These included functions related to aging hallmarks, including proteostasis, genomic maintenance, epigenetic regulation and autophagy. Several BP terms were enriched following both treatments (ESM Tables S7-S8).

Third, we investigated whether pharmacological lifespan extension is linked to reduced methylomic entropy. We assessed methylomic entropy in 35-day-old males treated with rapamycin or resveratrol, and compared it to age-matched controls. At the genome-wide level, neither treatment significantly altered methylomic entropy (mean methylomic entropy ± SD: 6mA: rapamycin = 0.32□±□0.01, resveratrol = 0.35□±□0.01, control = 0.34□±□0.03, Kruskal-Wallis: *chi-squared* = 5.956, *df* = 2, *p* = 0.51; 5mC: rapamycin = 0.36□±□0.02, resveratrol = 0.38□±□0.01, control = 0.33□±□0.01, one-way ANOVA: *F* = 1.06, *df* = 2, *p* = 0.54). However, when focusing on DMS, there was a significant decrease in 6mA entropy, but not 5mC entropy (6mA: rapamycin = 0.52□±□0.03, resveratrol = 0.51□±□0.03, control = 0.59□±□0.04, Kruskal-Wallis: *chi-squared* = 4.62, *df* = 2, *p* = 0.05; 5mC: rapamycin = 0.72□±□0.01, resveratrol = 0.73□±□0.01, control = 0.72□±□0.01, one-way ANOVA: *F* = 1.861, *df* = 2, *p* = 0.25).

Finally, we evaluated whether the different epigenetic clocks capture signals of biological aging by comparing the predicted epigenetic age of males treated with rapamycin or resveratrol to that of untreated control males of the same chronological age (35-day-old). Notably, all 6mA and 5mC clocks predicted that the epigenetic age of at least one treatment group (rapamycin- or resveratrol) was significantly lower than control males (Figures 3D), indicating that both rapamycin and resveratrol effectively decelerate the rate of epigenetic aging.

## Discussion

Epigenetic alterations that occur during aging are increasingly recognized as the gold standard proxy for both chronological and biological age across species. In this study, we explore the aging DNA methylome of an insect, the buff-tailed bumblebee *Bombus terrestris*, and compare age-related changes in N6-methyladenine (6mA), an epigenetic mark that has virtually never been studied in the context of aging, and C5-methycytosine (5mC), which is established as a reliable aging biomarker. We show that both methylation types undergo comparable age-related variations in global, sites-specific, entropy and ROC levels. Furthermore, using linear age-related changes, we developed various epigenetic clocks using 6mA or 5mC whose epigenetic age prediction were comparatively accurate. These findings underscore the potential of 6mA as a biomarker of aging.

Aging is commonly associated with a global decline in DNAm across the genome, accompanied by localized regions of hypermethylation, particularly at regulatory sites such as promoters (11, 31, 32). Yet, genome-wide levels of both 6mA and 5mC remain stable during aging in *B. terrestris* males. This stability suggests two possible, non-mutually exclusive explanations: either methylation marks are inherently stable during aging in this species, or the DNAm maintenance machinery operates with high fidelity. The hyper-mutability of 5mC (36), taken together with our findings that the aging DNA methylome of *B. terrestris* experience several site-level variations, suggests that the overall methylation landscape is tightly preserved due to high DNAm maintenance fidelity. This may be especially critical in species whose genome harbor low global levels of both 6mA and 5mC that must be maintained to avoid near-complete loss, with potential consequences for genome stability and gene regulation. Comparative studies of methyltransferase expression and enzymatic efficiency between species with high (*e*.*g*., mammals) *versus* low (*e*.*g*., insects) global methylation levels would help determining whether lowly-methylated species have evolved more efficient or more precise DNAm maintenance mechanisms to compensate for their limited epigenetic buffer.

In contrast with the overall stability of global methylation levels, site-specific changes occur within the aging DNA methylome of *B. terrestris* males. DMS for each methylation type are located in genes linked to distinct biological pathways. For instance, 6mA DMS were linked to developmental processes, reflecting the function of this methylation type in *Drosophila melanogaster* where it controls tissue-specific gene regulation during development (37). In contrast, 5mC DMS are linked to several pathways linked to proteostasis—a well characterized hallmark of aging (3). The enrichment of 6mA and 5mC in distinct biological functions suggests that 6mA and 5mC capture different, complementary features of aging. Recent studies show that 5mC clocks align with some, but not all, aging hallmarks (18, 38), highlighting the potential value of incorporating 6mA into multi-omic aging biomarkers. A small yet significant proportion of 6mA and 5mC DMS are located within non-coding ncRNA, which play key roles in epigenetic regulation and have been implicated in aging processes (39–41). Notably, some 6mA DMS—but none of the 5mC DMS—were found within rDNA. Previous work reports that epigenetic clocks based solely on rDNA can achieve comparable age prediction accuracy to clocks built from genome-wide data (42). Therefore, 6mA in rDNA represents a promising target for developing cost-effective epigenetic clocks using targeted sequencing approaches.

Entropy and ROC analyses revealed that both stochastic and consistent linear alterations occur in the *B. terrestris* DNA methylome during aging, albeit at relatively low magnitudes. These modest shifts are consistent with our earlier hypothesis that the DNAm maintenance machinery in this species operates with high fidelity, preserving epigenetic patterns over time. Notably, the average ROC values for both 6mA and 5mC were comparable to the mean per-site methylation change observed among the 353 CpG sites of the Horvath epigenetic clock (31), further supporting the idea that the pace of epigenetic aging may be relatively conserved across animal taxa (12).

Distinct features of the aging DNA methylome show that 6mA and 5mC are reliable markers of chronological age as both epigenetic clocks constructed from site-specific methylation and entropy levels provided accurate age predictions. A central goal of aging research is to develop biomarkers that not only track chronological age but also reflect biological age which better predicts inter-individual aging differences and intrinsic mortality risk. Using rapamycin and resveratrol, we significantly extended both *B. terrestris* males’ mean and maximum lifespan, further expanding the range of species whose longevity is positively influenced by these agents (43–46). At the DNA methylome level, both agents altered the methylation of a restricted number of adenines and a large number cytosines, including some occurring within genes involved in key lifespan-regulating pathways. These genes include those related to the agents proteins—target of rapamycin (TOR) for rapamycin and NAD+-dependent protein deacetylase sirtuin-1 (SIRT1) for resveratrol—which are highly conserved regulators of lifespan across numerous animal species (47–49). In addition, almost half of enriched BP terms were common to rapamycin- and resveratrol-fed males, indicating that these agents act upon similar biological pathways.

Furthermore, several biological processes were enriched during natural aging (comparing 7-day-old *vs* 35-day-old control males) and pharmacologically induced lifespan extension (comparing control 35-day-old males *vs* 35-day-old males treated with rapamycin or resveratrol), indicating that these agents may extend lifespan by modulating pathways that are altered during normal aging. These findings reinforce the growing pool of evidence that aging may result in part from the persistence of molecular, cellular, and physiological processes that are advantageous early in life but that become deleterious over time (47). Therefore, it seems likely that lifespan-extending interventions enhance longevity by acting upon the biological pathways that underlie the natural aging process. In line with this, based on the predictions of the 6mA and 5mC clocks, the methylomes of the 35-day-old rapamycin- or resveratrol-fed males reflected a younger epigenetic age, a methylation state like that of chronologically younger individuals. This demonstrates that epigenetic clocks based on 6mA and 5mC effectively capture signals of biological aging and can detect interventions that slow the aging process.

Altogether, our study provides a proof of concept that 6mA can be used to develop robust epigenetic clocks that accurately predict both chronological and biological age in *B. terrestris*, supporting 6mA as a promising biomarker of aging, with the potential to complement or even substitute 5mC. To fully establish the utility and broad applicability of 6mA-based clocks, future studies should expand datasets across a broader range of species, tissue types, and environments.

## Materials and Methods

### Model organism

The buff-tailed bumblebee (*Bombus terrestris*) was used as a model system because (*i*) both 6mA and 5mC occur at very low levels in insect genomes (21, 50), including that of *B. terrestris* (51), making this species ideal for studying both DNAm types simultaneously, (*ii*) its relatively small genome size (∼393 Mb) (52) makes this species suitable for cost-effective Oxford Nanopore Technologies (ONT) sequencing, and (*iii*) we previously showed that the mean and maximum lifespan of *B. terrestris* workers can be experimentally extended using a single topical application of the pharmacological hypomethylating agent RG108, indicating that DNAm (5mC) plays a role in lifespan regulation in this species (51).

*B. terrestris* males were obtained from Biobest (Westerlo, Belgium). A total of 150 males from 10 different colonies of origin were randomly assigned to 15 experimental microcolonies so that each contained 10 males. Microcolonies were maintained under standard laboratory conditions (red light, temperature = 27 ± 1°C, relative humidity = 50–60%) and were provided with *ad libitum* access to sugar syrup (Biogluc, Biobest, Westerlo, Belgium). Preliminary survival experiments revealed that mean male lifespan in the lab is 27.4 ± 7.1 days (maximum = 51 days, *n* = 791). Therefore, we used males that were 7, 21, and 35 days old in our subsequent molecular analyses to represent the entire lifespan of bumblebee adult males. Since epigenetic age has been shown to oscillates during the day (53), individuals were sampled at the same time of the day (10-11 AM) to prevent circadian-related bias. Individuals were randomly sampled across microcolonies, flash frozen in liquid nitrogen, and stored at -80°C until further processing. In total, 15 bumblebee males were sequenced across the three age groups (*n* = 5 per age group).

### Oxford Nanopore Technology sequencing

Genome wide, base-resolution DNA methylation levels were generated with long-reads ONT sequencing. High molecular weight genomic DNA (gDNA) was extracted from the thorax and legs of males 7, 21, or 35-day-old (*n* = 15) using an in-house SDS/proteinase K protocol. The head and abdomen were discarded to exclude pheromonal head glands and the content of the digestive tract, respectively. Briefly, frozen tissues were individually dry ground, suspended in an SDS/proteinase K solution, and left in suspension overnight. Next, we performed phenol– chloroform/chloroform purification followed by gDNA precipitation using ethanol and sodium acetate. gDNA integrity was assessed using agarose gel electrophoresis. qDNA quantity and absorbance ratios were measured using a Qubit 3.0 fluorometer (Thermo Fisher Scientific) and a NanoDrop ONE spectrophotometer (Thermo Fisher Scientific), respectively. gDNA libraries were prepared with the Ligation Sequencing gDNA - Native Barcoding Kit 96 V14 (ONT, SQK-NBD114.96) and sequenced with a PromethION platform (ONT, PRO-SEQ002).

### DNA methylation analyses

Raw reads were basecalled using Dorado (ONT, v. 0.7.2) with the super high accuracy model (‘sup’ command) to capture signals of modified bases (6mA and 5mC) in any genomic context. Basecalled reads aligned to the *B. terrestris* GCF_910591885.1 v. 1.2 reference genome (https://www.ncbi.nlm.nih.gov/datasets/genome/GCF_910591885.1/) using Dorado (‘aligner’ function). SAMtools was used for indexing, sorting and coverage quantification (mean coverage per sample ± SD = 32.6 ± 10.7X). Genome-wide global methylation levels were generated based on aligned BAM files using Modkit (ONT, v. 0.3.2) ‘summary’ command. Genome-wide, base-resolution methylation levels for 6mA and 5mC were generated using Modkit (ONT, v. 0.3.2) ‘pileup’ command. Only sites with a minimum coverage of 10X were kept in the subsequent analyses. After coverage filtering, 85,228,908 ± 19,234,608 adenines and 146,060,972 ± 32,906,853 cytosines were conserved. All subsequent analyses were conducted separately for 6mA and 5mC. Differential methylation analysis was performed between 7-day-old and 35-day-old males using Modkit (ONT, v0.3.2) ‘dmr pair’ function to identify significant age-related differentially methylated sites (DMS) for 6mA or 5mC. Functional enrichment analyses using gene ontology (GO) terms were conducted by linking DMS with biological processes (BP) GO terms from Hymenoptera Genome Database (54). Gene IDs were found at: www.ncbi.nlm.nih.gov/data-hub/gene/taxon/30195/).

Methylation entropy and ROC were calculated as described in (11). Briefly, site-level methylation entropy was quantified, then averaged to sample-level entropy. ROC analysis was performed by fitting simple linear models to age-related changes in 6mA or 5mC levels. Significant models (*p* < 0.05 and R^2^ > 0.5) were averaged to sample-level ROC.

### Epigenetic clocks

Epigenetic age prediction models (epigenetic clocks) were developed using penalized regression models to identify sites whose methylation tracks chronological age. To ensure model generalizability, we randomly split our initial dataset (*n* = 15) into training (*n* = 10) and testing (*n* = 5) datasets. Sample splitting between training/testing datasets was consistent between each model to ensure comparability. Three types of epigenetic clocks were developed for each DNAm type (6mA or 5mC): elastic net regressions based on 6mA or 5mC levels, ridge regression based on 6mA or 5mC levels of significantly DMS, and elastic net regressions based on 6mA or 5mC entropy levels. While elastic net regressions (alpha = 0.5) perform feature selection to only retain sites highly that correlate with chronological age, ridge regressions (alpha = 0) retain all input sites but shrinks the model coefficient of weakly predictive sites toward zero without driving them entirely to zero. For each model, a leave-one-out cross-validation (LOOCV) approach was implemented to perform model training independently within *n* fold (*i*.*e*., 15 folds), which reduces the risk of overfitting. The LOOCV was nested into outer loop for performance estimation and inner loop for lambda hyper-parameter fine-tuning. Predicted epigenetic age was calculated based on the combined methylation/entropy values of the selected adenines or cytosines, which were weighted by their respective coefficients.

### Pharmacological extension of bee lifespan

To test whether our epigenetic clocks track biological age, in addition to being highly predictive of chronological age, we sought to experimentally induce variations in biological age between individuals of the same chronological age. To this end, we used two well-studied pharmacological agents that increase lifespan in a wide range of species—rapamycin and resveratrol—to increase adult male lifespan and assess how they influence epigenetic age. We proceeded in two consecutive steps.

First, we compared the lifespan and survival of rapamycin- or resveratrol-treated *versus* control males to confirm whether these agents indeed prolong lifespan in *B. terrestris* males. Males were fed (*i*) sugar syrup solution containing DMSO (*n* = 49), which is the solvent used for rapamycin and resveratrol dilution, (*ii*) sugar syrup solution containing rapamycin (SelleckChem, S1039, final concentration = 100 μM, *n* = 32) —an inhibitor of mTOR, a protein complex that regulates growth, reproduction, and lifespan in diverse animal species (55), or (*iii*) resveratrol (MedChemExpress, HY-16561, final concentration = 100 μM, *n* = 31) —an activator of the longevity-promoting SIRT1 protein (56). Sugar syrup containing rapamycin or resveratrol was renewed three times per week to ensure optimal levels of drug activity. Treatment feeding started at 7-day-old and continued until male death. The effects of rapamycin and resveratrol on survival curves was compared between treatment groups using Cox proportional hazards model with mixed effects (*coxme* package in R (57)) in which experimental microcolony was computed as random factors.

Second, we tested if rapamycin and/or resveratrol affected the aging DNA methylome of *B. terrestris* males by sampling and sequencing 35-day-old males fed a control solution (sugar syrup), rapamycin or resveratrol (*n* = 3 per condition). DNA extraction, library preparation, ONT sequencing, differential methylation analysis, entropy quantification, and epigenetic age prediction were conducted as described for the initial dataset.

### Statistical Analysis

All statistical tests were performed in the R programming environment (version 2023.06.0+421 (2023.06.0+421), except for differential methylation analyses which were conducted with Modkit in the Bash environment. Modkit is a ready-to-implement tool from ONT that uses a Bayesian framework to estimate differential base modification (e.g., methylated *versus* unmethylated) between conditions. It uses a beta distribution to generate a posterior probability of modification at each site, then derive a Maximum A Posteriori (MAP)-based p-value which quantifies the confidence in observed differences. The MAP-based p-value considers effect size and coverage from Modkit pileup-generated files to ensure statistical robustness of site-resolution differential methylation detection (https://nanoporetech.github.io/modkit/dmr_scoring_details.html). False discovery rate (FDR) correction to control type I error was implemented using the Benjamini-Hochberg method. Sites were considered as statistically differentially methytlated (DMS) when FDR < 0.05.

Functional enrichment analyses were conducted using the R package topGO (58) with the ‘weight01’ algorithm, which considers terms dependencies when performing statistical analyses, thus reducing redundancy and, consequently, minimizing false positives while retaining biologically-relevant information.

Methylomic entropy was calculated by averaging mean per-site methylation entropy to provide a single entropy value per sample (11). Methylomic entropy was compared for 6mA and 5mC between the three age groups using one-way-ANOVA after confirming normality and homoscedasticity with Shapiro and Levene tests, respectively.

Rate of change (ROC) analyses were performed by fitting simple linear regression models to age-related changes in 6mA or 5mC DMS. The slopes were derived to quantify the percentage methylation change per unit of time (*i*.*e*., weeks). Sample-level mean ROC was calculated by averaging the slopes of statistically significantly fitted models (*p* < 0.05 and R^2^ > 0.65).

Epigenetic clocks were developed using penalized regression models. Briefly, samples were split into training (*n* = 10) and testing (*n* = 5) datasets with proportions roughly equal to 65% and 35% of the total dataset (*n* = 15). Given our limited sample size, we chose to remove sites missing methylation values, rather than applying imputation methods, to minimize the risk of overfitting resulting from introducing artificial artefacts. The glmnet R package (59) was used to regress methylation matrices against known ages in the training set, then applying the model to the naïve (testing) dataset to assess model performances. Elastic net and ridge regressions were implemented by setting the alpha hyperparameter to 0.5 or 0, respectively. Mean absolute error (MAE), mean relative error (MRE) and Pearson’s correlation (corr) were used as model performance estimation metrics.

### Data and material availability

All data generated or analysed during this study are included in this published article and its electronic supplementary material. ONT sequencing data are accessible in the NCBI SRA database.

## Acknowledgments

This work was supported by funding from the Belgian Fund for Scientific Research (FRS-FNRS), in the form of grants #C/RFE24/0071 (to TR), FC 57457 (to MB), and PDR T.0010.24, and by funding from the *Action de Recherches Concertées* – ULB in the form of grant #ARC 2025-2028 (to SA).

